# Cryo-EM analysis of human mitochondrial Hsp90 in multiple tetrameric states

**DOI:** 10.1101/2020.11.04.368837

**Authors:** Yanxin Liu, Ming Sun, Daniel Elnatan, Adam G. Larson, David A. Agard

## Abstract

Hsp90 is a ubiquitous molecular chaperone that mediates the folding and maturation of hundreds of “client” proteins. Although Hsp90s generally function as homodimers, recent discoveries suggested that the mitochondrion specific Hsp90 (TRAP1) also forms functionally relevant tetramers. The structural mechanism of tetramer formation remains elusive. Here we used a combination of solution, biochemical and cryo-electron microscopy (cryo-EM) approaches to confirm that, independent of nucleotide state, a subpopulation of TRAP1 exists as tetramers. Unexpectedly, cryo-EM reveals multiple tetramer conformations having TRAP1 dimers arranged in orthogonal, parallel, or antiparallel configurations. The cryo-EM structure of one of the orthogonal tetrameric states was determined at 3.5 Å resolution. Each of the two TRAP1 dimers is in a symmetric AMP·PNP-bound closed state with the tetramer being stabilized through three distinct dimer-dimer interaction sites. In unique ways, each of the three TRAP1 domains contributes to tetramer formation. In addition to tetramerization via direct dimer-dimer contacts, our structure suggests that additional stabilization could come from domain swapping between the dimers. These results expand our understanding of TRAP1 biology beyond the conventional view of a functional dimer and provide a platform to further explore the function and regulation of tetrameric TRAP1 in mitochondria.

## Introduction

Maintaining the proteome integrity is an essential function for all organisms. Key to this process are molecular chaperones such as Hsp90 that mediate the folding and maturation of hundreds of “client” proteins^1,2^. Higher eukaryotes possess four isoforms of Hsp90^3^: two in the cytosol (Hsp90α and Hsp90β), one in the endoplasmic reticulum (Grp94), and one in mitochondria (TRAP1). Hsp90s have a highly conserved modular structure with a nucleotide-binding N-terminal domain (NTD), a client-interacting middle domain (MD) and C-terminal domain (CTD)^4^. It is well established that Hsp90 functions as a homodimer with the CTD contributing to initial dimerization^5–13^. However, a recent discovery suggests the mitochondrial specific TRAP1 also functions as a tetramer^14^. The tetramer population correlates with the state of mitochondrial oxidative phosphorylation and appears to play a key role in mitochondrial metabolic homeostasis^14^. Despite the importance of the TRAP1 tetramer for mitochondrial function, the architecture of TRAP1 tetramer and how TRAP1 arranges its domains to form the tetramer (presumably via dimer-dimer interactions) remain elusive.

Here we employed orthogonal solution-based approaches, including blue native gel electrophoresis, analytical ultracentrifugation, and cryo-electron microscopy (cryo-EM), to confirm that recombinant human TRAP1 exists as an equilibrium between dimer and tetramer. Tetramer formation was not observed for either the bacterial or yeast cytosolic Hsp90s. Surprisingly, cryo-EM reveals that at least four different tetrameric states exist for closed state of TRAP1, in which the dimers are arranged in orthogonal, parallel, or antiparallel configurations. By designing a covalently linked TRAP1 heterodimer construct using SpyTag and SpyCatcher and fused with its client protein succinate dehydrogenase B (SdhB)^12,15,16^, we determined the cryo-EM structure for one of the orthogonally arranged TRAP1 tetramers at 3.5 Å resolution revealing the structural mechanism underlying tetramer formation. Our structure also suggests how domain swapping between TRAP1 dimers could further stabilize the tetramer.

## Results

### TRAP1 tetramer formation is nucleotide independent

Previous immunoblot results showed that both endogenous and recombinant human TRAP1 exist as an equilibrium between dimers and tetramers^14^. We confirmed this result using blue native PAGE and further explored to what extent tetramer formation depends on nucleotide binding. We ruled out the possibility that non-specific disulfide bonds were responsible for the tetramer formation, as the migration pattern remained unchanged in the presence of 100 mM of the reducing agent DTT (Fig. 1A). Surprisingly, TRAP1 tetramer formation was also independent of incubation with non-hydrolyzable ATP analog AMP•PNP (Fig. 1A). It is well appreciated that binding of non-hydrolyzable or slowly hydrolyzable ATP analogs universally induce a large conformational switch from an open to a closed state in all tested Hsp90s^4^. This is evidenced here as the AMP·PNP-bound TRAP1 tetramer ran as a narrower and faster-migrating band in blue native PAGE (Fig. 1A). A tetramer was not observed for either yeast cytosolic Hsc82 or bacterial Hsp90 (HtpG) under the same conditions (Fig. S1) suggesting this is a unique property of the mitochondrial Hsp90. The existence of a TRAP1 tetramer was further confirmed using analytical ultracentrifugation (AUC). The TRAP1 dimer and tetramer in the apo state were observed as distinct peaks with sedimentation coefficients of 9.1 S and 13.3 S, respectively (Fig. 1B). The ratio between dimer and tetramer population is 7:1, consistent with the intensities of corresponding bands from the blue native gel (Fig. 1A).

**Figure 1.**
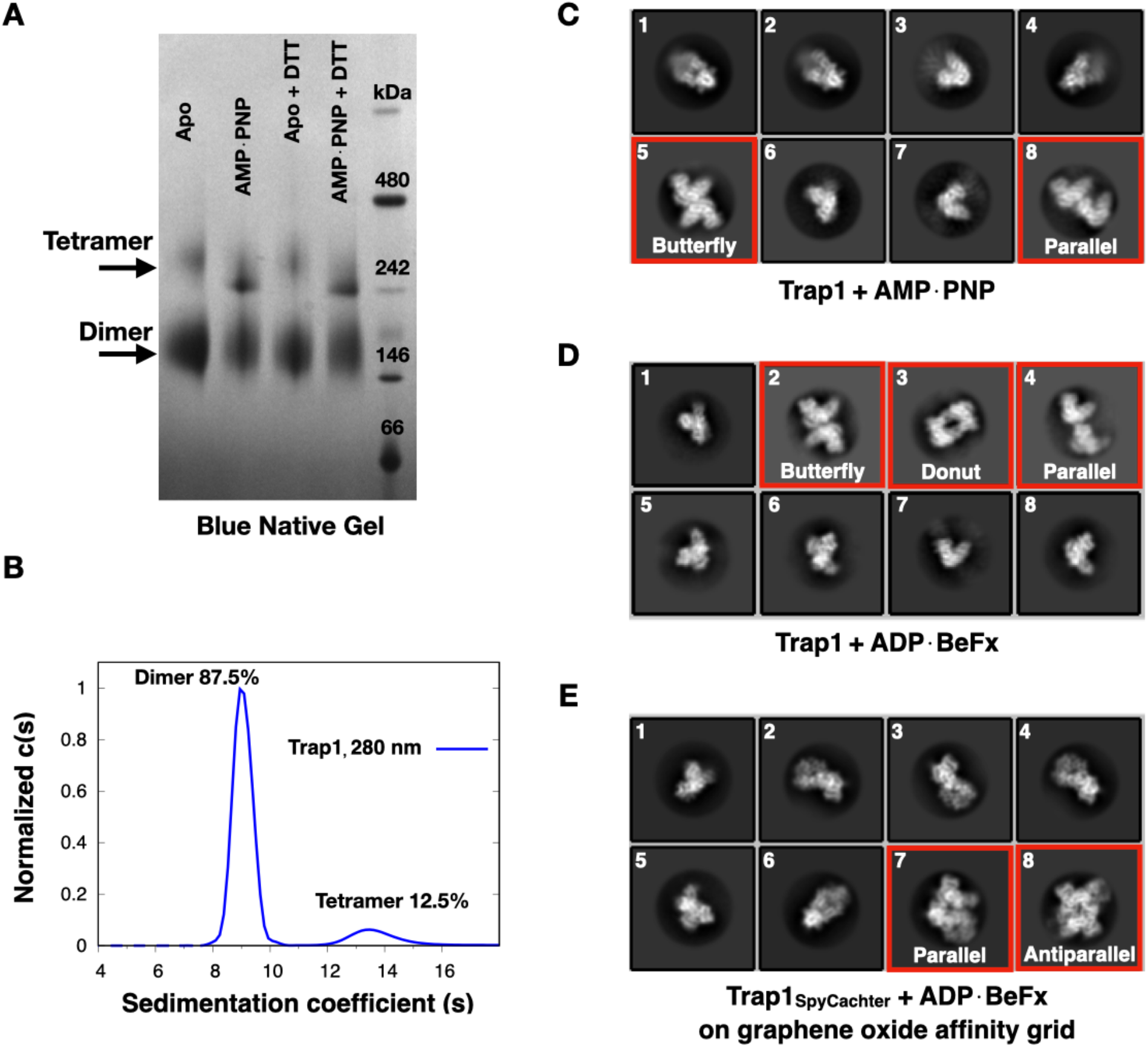
The equilibrium between dimer and tetramer population for recombinant human TRAP1. (A) Purified TRAP1 was analyzed by blue native gel electrophoresis and stained with Coomassie blue. The existence of TRAP1 tetramer is independent of nucleotide and reducing agent. (B) The equilibrium between TRAP1 dimer and tetramer was analyzed by sedimentation velocity analytical ultracentrifugation. *c*(*S*) is the sedimentation coefficient distribution. (C-D) Representative 2D-averaged classes of TRAP1 in the presence of AMP·PNP or ADP·BeF_x_ from single particle cryo-EM. Tetramers are boxed in red squares. (E) Representative 2D-averaged classes of ADP·BeF4_x_ bound TRAP1 fused with SpyCatcher at the C-terminus (TRAP1_SpyCatcher_). The SpyCatcher was used to covalently link TRAP1 to SpyTag on the surface of the graphene oxide affinity grid^17^. Tetramers are boxed in red squares. All 2D-averaged classes were grouped into four classes based on their shape or orientation: butterfly, donut, parallel, and antiparallel.

### Cryo-EM reveals four classes of TRAP1 tetramer

We had previously observed the TRAP1 tetramer in the presence of AMP·PNP using single-particle Cryo-EM^14^. However, despite significant efforts at optimization, only a single view could be observed, preventing us from obtaining a 3D structure. Here we collected a much larger dataset (the number of micrographs increased from 650 to 4500) in the hope of capturing more angular views. In addition to the previously observed “butterfly” tetramer conformation (class 5 in Fig. 1C), a second tetramer was obtained (class 8 in Fig. 1C) having a parallel arrangement of the two TRAP1 dimers. This is different form the orthogonal “butterfly” conformation and must therefore be using a different dimer-dimer interface. Changing nucleotide from AMP·PNP to ADP·BeF_x_ did not improve the angular distribution of the tetramer classes (Fig. 1D). Interestingly, this did result in the appearance of a third tetrameric “donut” shaped conformational class (class 3 in Fig. 1D). Adding to the complexity, we had previously observed two TRAP1 tetramer classes when using graphene oxide affinity grids to determine the cryo-EM structure for human TRAP1^17^ (class 7 and 8 in Fig. 1E). These two classes represent TRAP1 side views, having parallel and antiparallel arrangements, respectively. In summary, there are at least four distinct TRAP1 tetrameric classes. According to their shape or orientation, we classified them as butterfly, donut, parallel, and antiparallel classes.

Based on our knowledge of TRAP1 dimer structures and their 2D projections, we were able to arrange the dimers to form tetramers that matched the observed 2D-averaged projections. The “butterfly” tetramer corresponded to two closed TRAP1 dimers aligned orthogonally (Fig. 2A). The “donut” class in Fig. 1D corresponded to a second orthogonally arranged dimer of dimers, where the C/D dimer flips 180° relative to “butterfly” tetramer (Fig. 2B). The “parallel” tetramer was captured in two different views (Fig. 3A). Class 7 in Fig. 1E represented the side view of two closed TRAP1 dimers, whereas class 8 in Fig. 1C and class 4 in Fig. 1D were top views. We only observed side views for the “antiparallel” class 8 in Fig. 1E, which resembles the packing found in the zebra fish TRAP1 crystal structures (Fig. 3B)^8^. Although we can determine the spatial arrangement of dimers in all tetramer classes, the very restricted orientations precluded obtaining true 3D reconstructions for any of them. Therefore, the detailed nature of the dimer-dimer interface was unknown.

**Figure 2.**
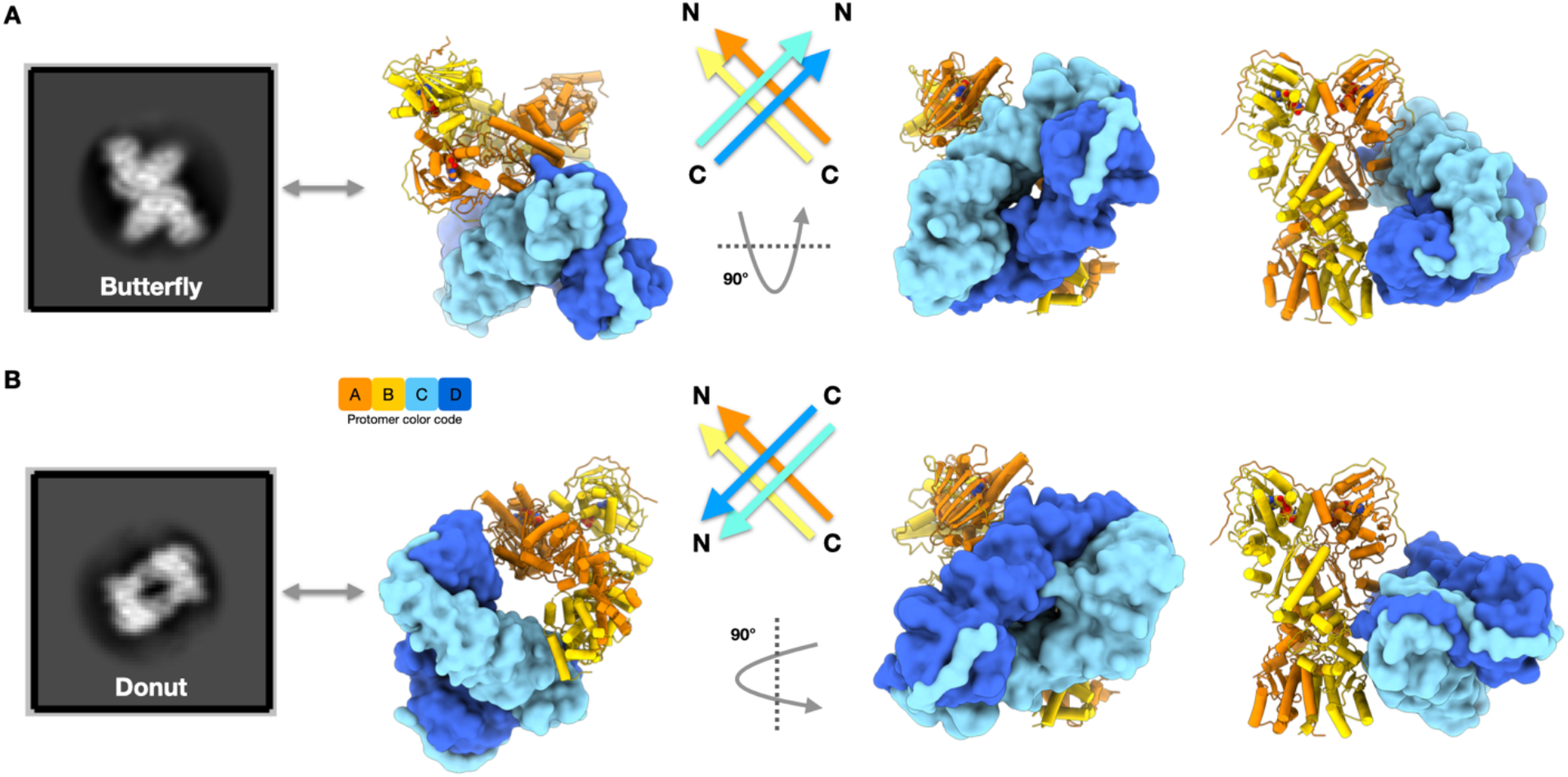
Structural modeling for the “butterfly” and “donut” classes of TRAP1 tetramer by orthogonally arranging two TRAP1 dimers in nucleotide-bound closed state. (A) Comparison between the “butterfly” tetramer conformation observed in cryo-EM and the structural model in different views. (B) An alternative orthogonally arranged dimer of dimers matches with the “donut” class of TRAP1 tetramer. It differs from the “butterfly” tetramer by a 180° flip of C/D dimer relative to A/B dimer.

**Figure 3.**
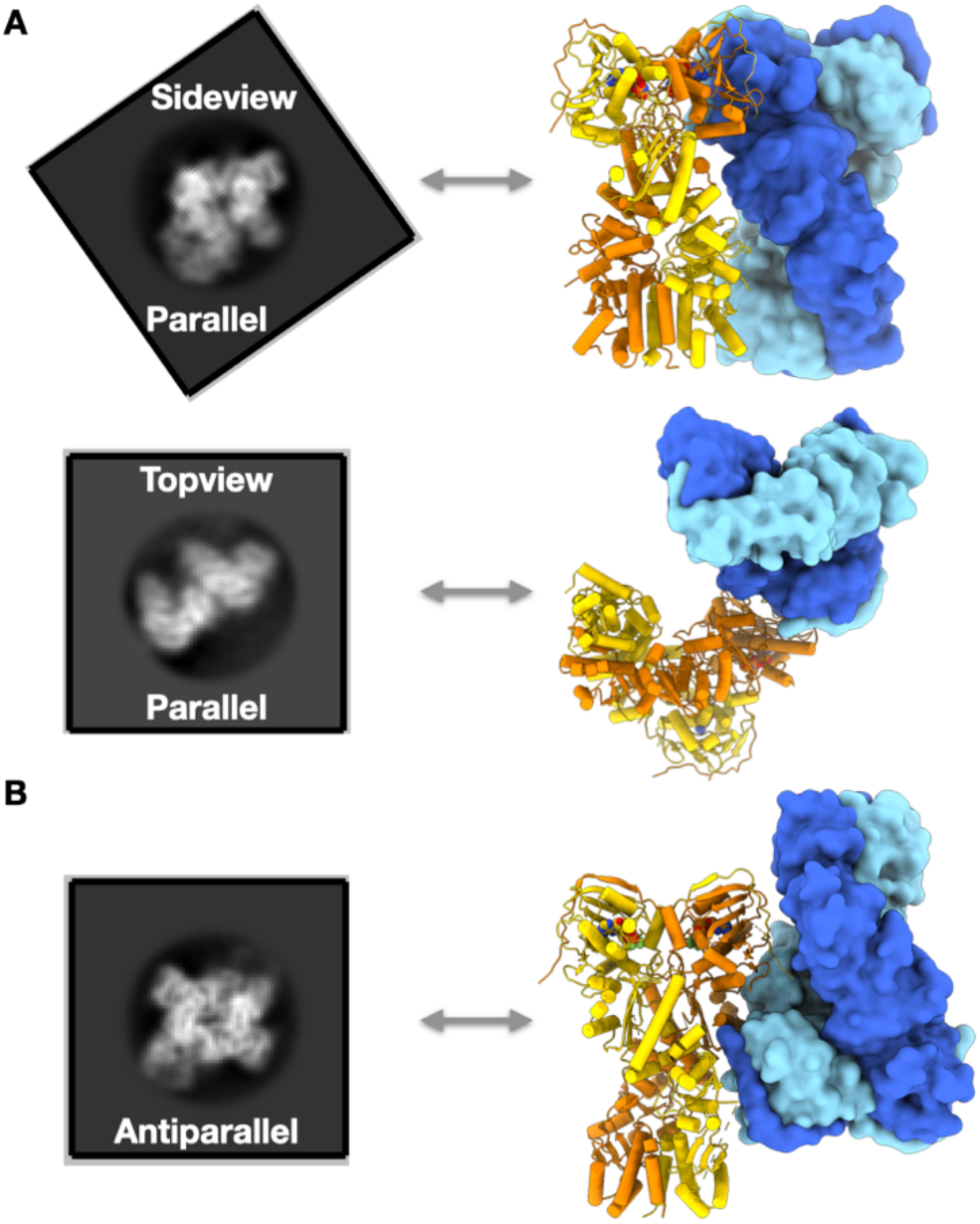
Structural modeling for the “parallel” and “antiparallel” classes of TRAP1 tetramer. (A) Comparison between the “parallel” tetramer observed in cryo-EM and the structural model in two different views. (B) Comparison between the “antiparallel” tetramer observed in cryo-EM and the packing in the zebra fish TRAP1 crystal structures^8^.

### Cryo-EM structure of a TRAP1 tetramer

To improve the TRAP1 dimer stability and potentially the tetramer angular distribution, we engineered a covalently linked TRAP1 heterodimer using SpyTag and SpyCacther^18,19^, referred to as construct TRAP1_SpyTag-SpyCacther_, to stabilize TRAP1-CTD dimerization. Although this construct improved the quality of particles in the TRAP1 dimer classes, it failed to improve the angular distribution of the tetramer. Using either AMP·PNP or ADP·BeF_x_, only a single view of the “butterfly” tetramer conformation was present (Fig. S2).

We have recently shown that the binding of the TRAP1 client protein SdhB further stabilized the TRAP1 dimer and induced an unexpected conversion of the intrinsic asymmetric closed state to a symmetric closed state^12^. Trying to combine both benefits, we fused SdhB after the SpyTag in our TRAP1_SpyTag_ construct, resulting in a covalently linked TRAP1 heterodimer with SdhB attached to one protomer (Fig. 4A), referred to as TRAP1_SdhB_. Consistent with having an attached client, the ATP hydrolysis rate for TRAP1_SdhB_ was ~2.5 faster than wild type TRAP1 and the TRAP1_SpyTag-SpyCacther_ heterodimer but was slower than wild type TRAP1 stimulated by saturating SdhB (Fig. S3). The accelerated ATPase activity of TRAP1_SdhB_ indicated that the fused SdhB must be interacting productively with TRAP1. Using TRAP1_SdhB_ in Cryo-EM, we again observed the “butterfly” conformation in 2D-averaged classes (class 4 in Fig. 4B), but now found previously unseen views of the tetramer (class 5 and 7 in Fig. 4B). From this dataset, we determined 3D structures of TRAP1 in dimeric and tetrameric states and refined them to 3.1 Å and 3.5 Å resolution, respectively (Fig. 4C-D and Fig. S4).

**Figure 4.**
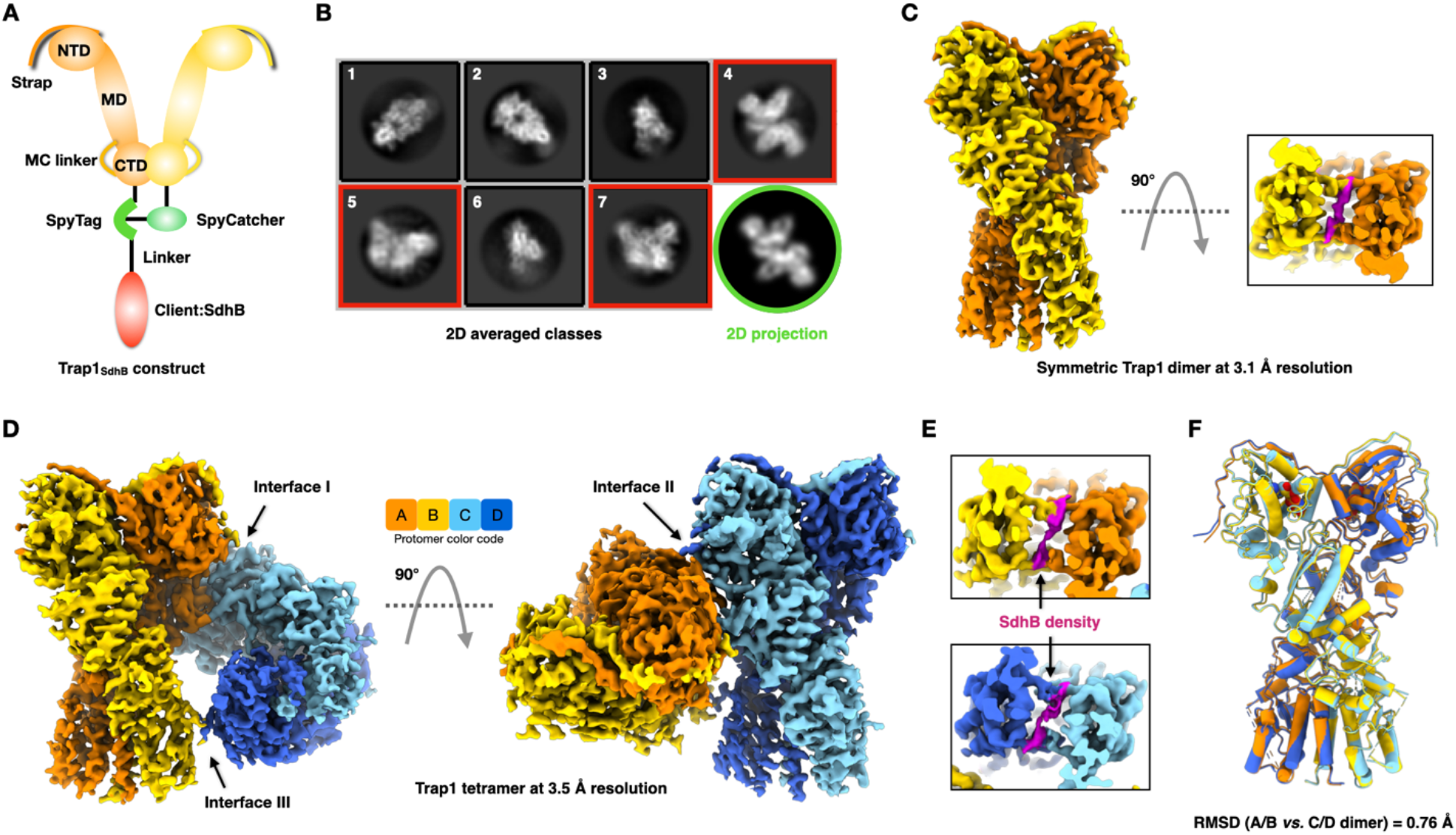
Cryo-EM structure of TRAP1 heterodimer in tetrameric and dimeric states. (A) The construct design of TRAP1_SdhB_. A client protein SdhB is fused to the TRAP1 heterodimer covalently linked with SpyTag and SpyCatcher. (B) Representative 2D-averaged classes of AMP·PNP-bound TRAP1_SdhB_ from single particle cryo-EM. The tetramers are boxed in red squares. The 2D projection of the final 3D reconstructed TRAP1_SdhB_ tetramer after low-pass filter at 10 Å was shown in green circle for comparison with the “butterfly” conformation in class 4. (C) The cryo-EM structure of TRAP1_SdhB_ in a dimeric state at 3.1 Å resolution. The SdhB density in the lumen is shown in magenta. (D) The cryo-EM structure of TRAP1_SdhB_ in a tetrameric state at 3.5 Å resolution. (E) The SdhB density in the lumens of both dimers in tetramer shown in magenta. (F) The structural alignment between TRAP1 A/B and C/D dimers in the tetramer (RMSD= 0.76 Å).

The dimeric TRAP1 is in a symmetric closed state with sparse density present in the TRAP1 lumen (Fig. 4C). This is similar to the TRAP1 structures we solved before in which free SdhB was bound to TRAP1. Client binding induced an asymmetric to symmetric transition in the closed state of TRAP1 (Fig. 5S)^12^. Despite being covalently linked to TRAP1 and contrasting with previous work, no density was observed for the SdhB-NTD globular domain. While covalent linkage may represent a general strategy for determining weakly bound protein complexes, owing to its similarity to our previous structure^12^, SdhB interactions will not be discussed here.

**Figure 5.**
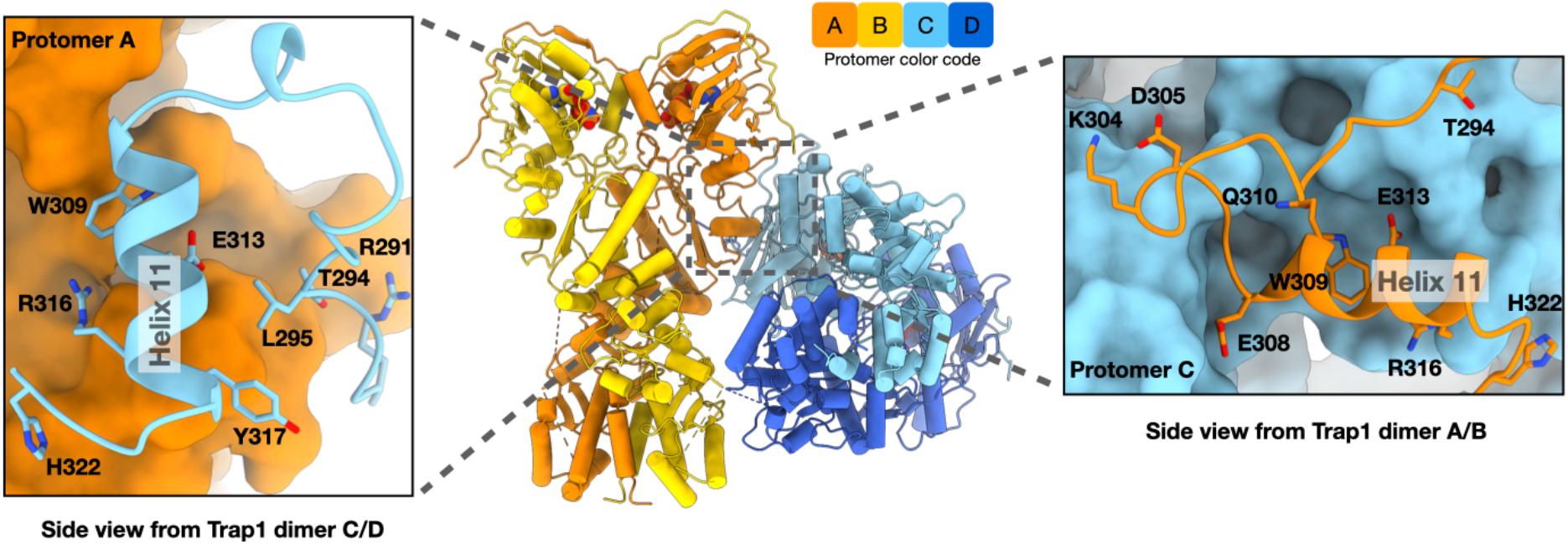
The MD-MD dimer-dimer interface I between protomer A and C. Inset on the left: the interacting residues from protomer C (light blue) at the interface of two TRAP1 dimers. The protomer A is depicted as orange surface. Inset on the right: The interacting residues from protomer A (orange) at the interface of two TRAP1 dimers. The protomer C is depicted as light blue surface.

As can be seen from 2D projections, the 3.5 Å tetrameric TRAP1 structure determined here corresponds to the “butterfly” class discussed above (Fig. 4B). This ruled out the possibility that the TRAP1 tetramer structure we obtained was somehow an artifact of the fused SdhB. None of the 2D projections matched with the other tetrameric 2D averaged classes of TRAP1 highlighted in Fig. 2 and 3, confirming that there are indeed multiple closed tetrameric states. In this “butterfly” tetramer class, two TRAP1 dimers (referred to as the A/B and C/D dimers) interact orthogonally to form a dimer of dimers (Fig. 4D). This contrasts markedly with the anti-parallel packing observed in the zebra fish TRAP1 crystal structure (Fig. 3B)^8,19^. Due to the presence of AMP·PNP, both dimers adopted a closed state with their NTDs fully dimerized. Continuous density corresponding to SdhB in the lumen can be seen in both dimers but could not be uniquely modeled due to the limited resolution (Fig. 4E). Similar to the dimeric structure, the conformation of each dimer within the tetramer closely resembled the SdhB-bound symmetric closed state observed for the non-covalent complex (Fig. S5)^12^, rather than the asymmetric closed state in the absence of client protein^8,17^. The symmetric TRAP1 state was at least partially induced by the fused SdhB and may be further stabilized by tetramer formation. The conformational difference between the A/B and C/D dimers was minimal (RMSD=0.76 Å; Fig. 4F). However, docking the C/D dimer into the A/B dimer density resulted in a significant mismatch (Fig. S6), indicating that the tetramer itself was asymmetric. Docking of two asymmetric closed TRAP1 dimers into the tetramer density resulted in various degrees of clashes (Fig. S7). Thus, the “butterfly” tetramer conformation observed in wild type TRAP1 without SdhB (Fig. 1C and 1D) must utilize a symmetric closed state or an asymmetric closed state distinct from any observed so far^8,12,17^.

### TRAP1 dimer-dimer interacting regions in middle domain

There are three interacting regions at the dimer-dimer interface that stabilize the TRAP1 tetramer. Interface I centered around helix 11 located at beginning of the MD domain in protomer A, which interacted with a similar but not identical region in protomer C (buried surface area of 581 Å^2^). Key residues at this interface are shown in Fig. 5. W309 from protomer A (W309_A_) formed a hydrophobic interaction with L295_C_ and Y317_C_. Because the dimers are perpendicular, W309_C_ formed a distinct hydrophobic interaction with T294_A_. Interestingly, W309 is not conserved among Hsp90s (Fig. S8). It is mostly hydrophobic in TRAP1s (Trp or Phe), whereas it is acidic (Asp or Glu) in all other Hsp90 homologs (except Asn in Hsp90C in chloroplast). At the core of this region is a salt bridge formed between E313_A_ and R316_C_. This interface is also stabilized by a network of hydrogen bonds between R316_A_, H322_A_, and H322_C_. TRAP1 H322 is only conserved among certain mammals. TRAP1s in rodent and lower species have a hydrophobic residue at this position, as found in all other Hsp90 homologs including the bacterial HtpG (Fig. S8). For cytosolic Hsp90s we have shown that the residue at this position can interact with client proteins^13^ or with cochaperones (such as Aha1)^11^ through hydrophobic interactions.

### The N-terminal straps form *trans*-dimer interactions with the middle domain

Interface II involves the N-terminal extension or strap in TRAP1^20^. While missing in bacterial and yeast Hsp90, the strap is found in cytosolic and organellar Hsp90s in most eukaryotes and forms *trans*-protomer interactions that stabilize the Hsp90 closed state^8,10^. We have previously identified the strap as a regulatory element that is responsible for the unique temperature dependence of TRAP1 ATPase activity^20^. Here in the tetrameric state, sparse strap density extends beyond the crystallographically resolved P70, and corresponds to the first 10 residues (60-69) of the matured human TRAP1 (Fig. 6A). However, the resolution of the strap density in this region is limited and was only visible at low contour level, suggesting flexibility and heterogeneity. This strap density contacts the *trans*-dimer MD which was well resolved with clear sidechain density. The residues involve in the strap-MD interaction are R421 and K424 in the Helix 13 and nearby Y331 (Fig. 6B). Together they form a positively charged patch on the MD surface that could electrostatically complement negatively charged residues in the strap (65-EDKEE-69) (Fig. 6C). Intriguingly, these charged strap residues and their contact on the MD are highly conserved in the TRAP1 family (Fig. S9). Furthermore, K424 has been identified as an acetylation site^21^ and Y331 is predicted to be a phosphorylation site^22^. Through posttranslational modifications, both residues could regulate the strap-MD interaction and consequently tetramer formation.

**Figure 6.**
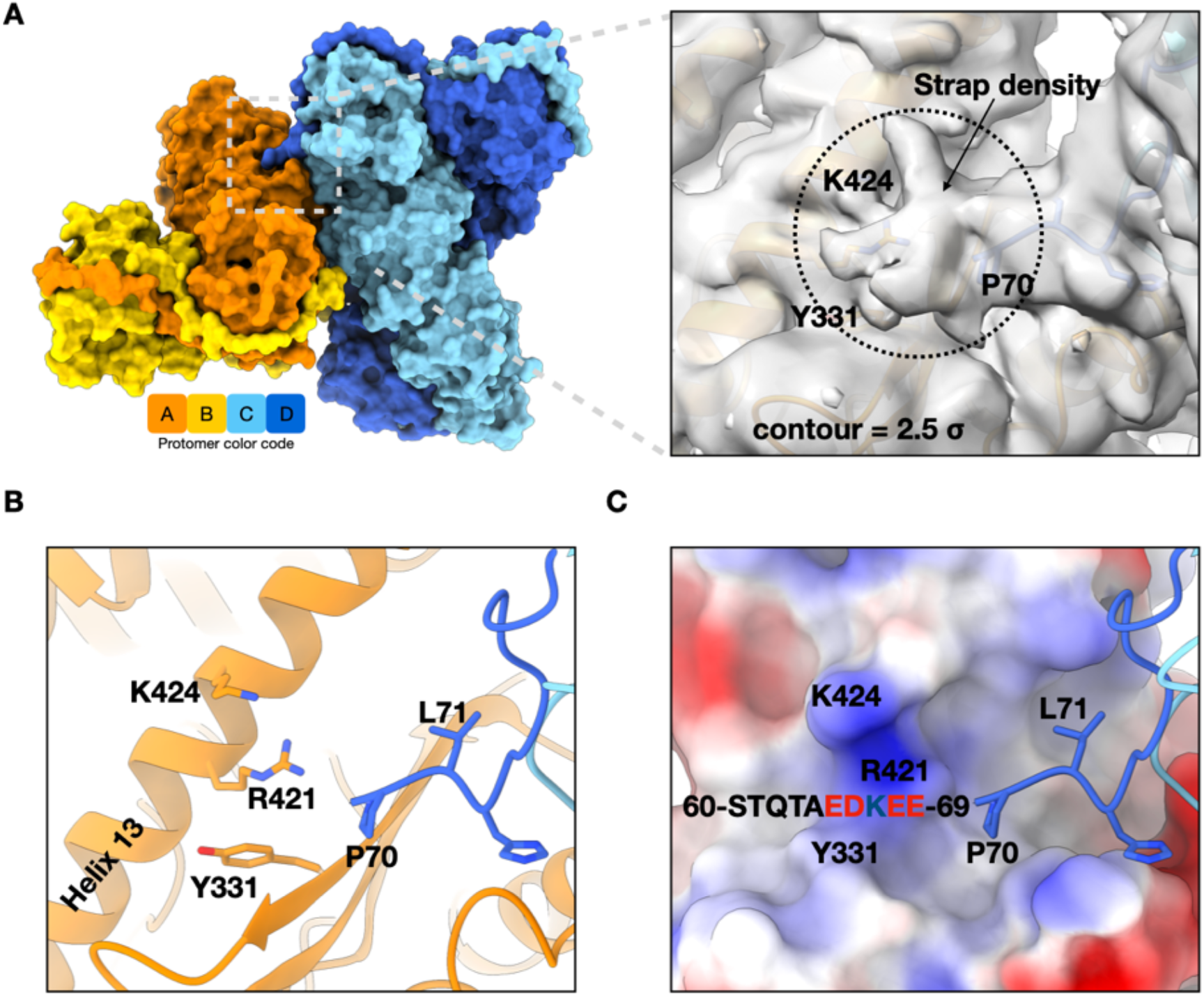
The strap-MD dimer-dimer interface II between protomer A and D. (A) Surface representation of the TRAP1 tetramer. Inset: a sparse strap density at low contour level corresponds to the first 10 residues (60-69) in the matured form of human TRAP1. (B) The residues involved in the strap-MD interaction are R421 and K424 in the Helix 13 and nearby Y331. (C) A positively charged patch on the electrostatic potential surface at the strap-MD interface, which is complementary to the negatively charged residues in the strap (65-EDKEE-69).

### The MC linker *trans*-dimer interaction and CTD domain swapping

Interface III involves the first few residues of the linker between the MD and CTD, known as the MC linker, from protomers B and D (Fig. 7A). Notably, this is a different pair of protomers than the ones that define Interface I (A and C in Fig. 5) and II (A and D in Fig. 6). Together, multiple contact sites across multiple pairs of protomers significantly stabilize the tetrameric TRAP1.

**Figure 7.**
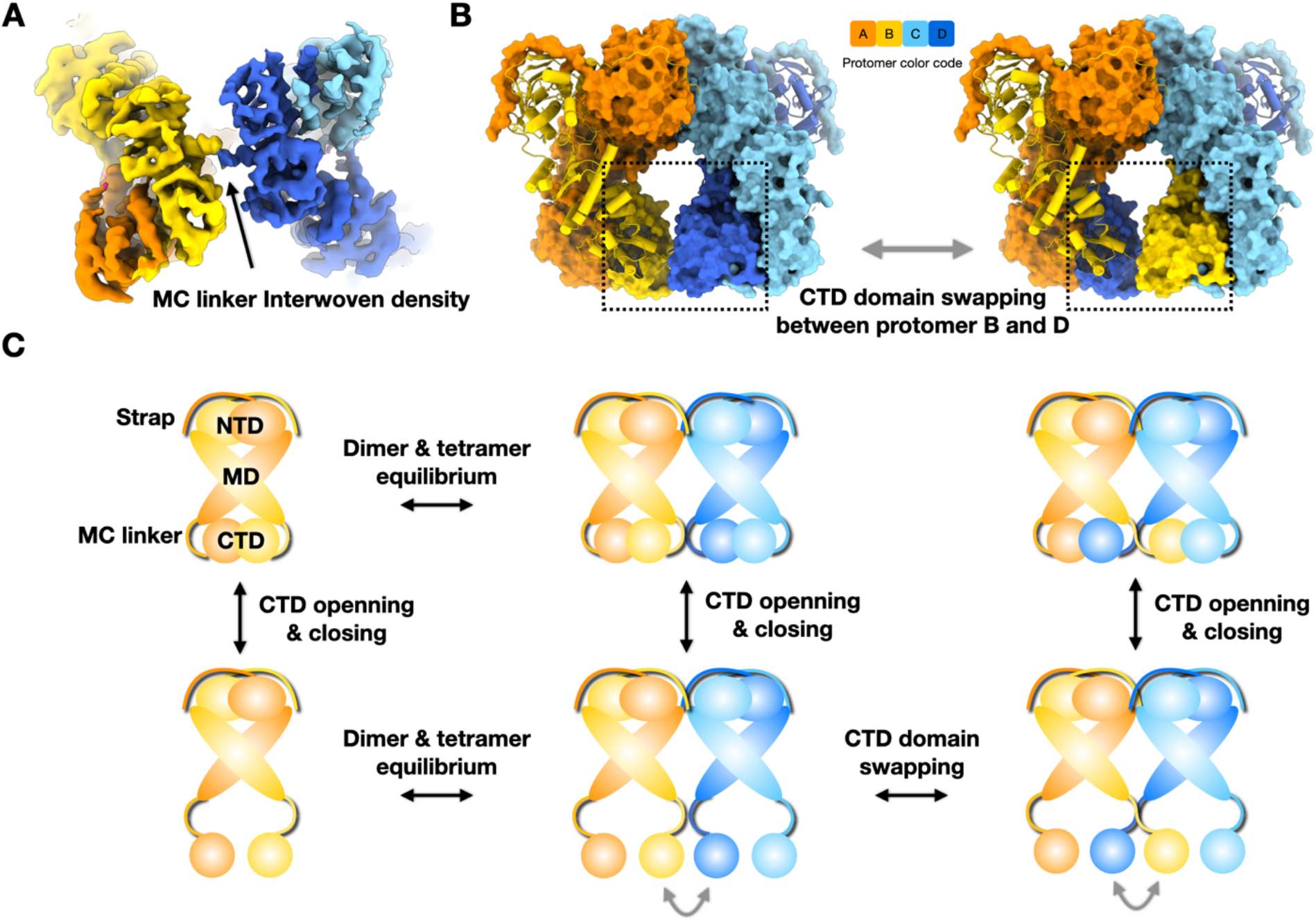
The MC linker dimer-dimer interface III suggests a model for CTD domain swapping. (A) The interwoven density between MC linkers of protomer B and D. (B) CTD domain swapping between B and D results in a more stable tetrameric state. (C) A tentative model for TRAP1 tetramer formation through CTD domain swapping. The nucleotide bound NTD-dimerized TRAP1 exist as an equilibrium between CTD open and closed states. Using the CTD closing and opening mechanism, TRAP1 could switch between two tetrameric states by CTD domain swapping.

The MC linker (558-HYKEEKFEDRSPAAE-572) is highly charged and is disordered in both the crystal structure and the cryo-EM structure of dimeric TRAP1^8,17^. Unfortunately, the limited resolution in this region prevented the building a reliable atomic model around this site. This region in protomers B and D could be stabilized through self-complementary electrostatic interactions. Considering the flexibility of the MC linker, an attractive possibility is that domain swapping occurs between the CTD domains of protomers B and D (Fig. 7B). Thus, the CTD from protomer B would replace the CTD from protomer D in the CD dimer and the protomer D-CTD would invade the AB dimer. If that were to occur, it would provide significant additional stabilization to the tetramer.

### A tentative model for TRAP1 tetramer formation

Owing to the disordered connector, the two tetramer models shown in Fig. 7B are indistinguishable by cryo-EM. Given the high likelihood that the CTDs within a wild type closed state can dynamically open, CTD domain swapping could occur easily (Fig 7C). We have consistently observed TRAP1 dimers with missing CTD density in cryo-EM structures of human TRAP1^12,17^ despite full-length protein being used. This is even observed when the CTDs are indirectly linked through SpyCatcher-SpyTag (Fig. S2 and S4).

Although CTD opening appears to be worsened by interactions with the air-water interface during sample freezing for cryo-EM, it still readily occurs even when TRAP1 is anchored away from the interface by using affinity grids^17^. Moreover, CTD opening has also been observed in the closed state of yeast Hsp90 by single-molecule FRET^23^, suggesting it is a common propensity with Hsp90s. Based on these observations, we propose a model for TRAP1 tetramer formation through CTD domain swapping (Fig. 7C). The CTD can open and close in both dimeric and tetrameric TRAP1, as long as TRAP1 is in the nucleotide bound closed state. Using the CTD opening and closing mechanism, TRAP1 could switch between two tetrameric states by CTD domain swapping. However, further experimentation is still needed to determine if domain swapping truly occurs.

## Discussion

A recent study established a link between TRAP1 tetramer formation and mitochondrial oxidative phosphorylation^14^. Here we report that TRAP1 can form at least four distinct tetramer configurations in the nucleotide-bound closed state. Solving the structure of one of these states required the use of a specially designed covalently linked TRAP1 heterodimer with one protomer fused to the TRAP1 client, SdhB. The fact that the same “butterfly” tetramer readily forms with wild type TRAP1 in the absence of SdhB or SpyCatcher/SpyTag ruled out the possibility of it being an artifact. From the 3D structure, three *trans*-dimer interfaces were identified. Unexpectedly, each interface utilized a different pair of protomers.

In addition to the TRAP1 tetramer in the closed state, blue native gels and AUC showed that TRAP1 also forms a tetramer in the open apo state. Despite extensive efforts we were not able to visualize the open TRAP1 tetramer in cryo-EM presumably due to its flexibility. Docking of the HtpG open V-shape structures into our TRAP1 tetrameric state suggests that the main MDMD interface I could be utilized by apo TRAP1 dimers, but steric clashes would require that two of the TRAP1-NTDs be undocking from the TRAP1-MD (Fig. S10). What makes this plausible is that the equivalent undocking has been observed to occur spontaneously in yeast Hsp90 using optical tweezers^24^. Similarly, binding of the Aha1 cochaperone also induces NTD undocking in apo yeast Hsp90^11^. Thus, tetramer formation in TRAP1 may provide a similar means to accelerate closure and regulate the TRAP1 conformational cycle without the need for a special cochaperone.

We recently *in vitro* reconstituted SdhB as a model TRAP1 client and provided the first insights into TRAP1-client protein interactions by determining the cryo-EM structure of TRAP1-SdhB complex. Surprisingly, in the presence of client, the normal asymmetric TRAP1 closed state became symmetric^12^. In the current work, the presence of SdhB fused to the C-terminus of one TRAP1 protomer also drove formation of the symmetric closed state, but only SdhB density within the TRAP1 lumen was observed. Important for thinking about tetramer function, the large non-lumenal SdhB N-terminal region in the complex^12^ could be accommodated, but only if it were bound to the outer face of either dimer (Fig. S11). Thus, while the symmetry conversion seems intrinsic to having a client bound, there seems to be a complex relationship between client and tetramer formation. Furthermore, no tetramers were visible in our previous work where TRAP1 dimer bound with free SdhB^12^. This result suggests that each oligomerization (dimer or tetramer) state could serve a different subset of client proteins.

In our tetramer structure, client density was observed in the lumens of both compositional dimers, indicating that both are functional for client binding. This raises the interesting possibility that the TRAP1 tetramer could bind multiple parts of a single client protein, stabilizing a much more extended conformation or acting on different domains. This would be somewhat analogous to how multiple Hsp70s can bind to one client protein^25,26^, but with the benefit of being more precessive. Alternatively, the two TRAP1 lumens could be used by different client proteins, perhaps facilitating complex assembly.

Our structure of the TRAP1 tetramer contributes significantly towards understanding the unique properties of TRAP1 and how it regulates mitochondrial metabolism through its interaction with client proteins^15,16,27^. Moving forward, the high-resolution structural determination of other TRAP1 tetramers states in both closed and open states remain important future research directions. These will validate our hypothetical models based only on cryo-EM 2D averages of the tetrameric states. What also remains to be explored is how mitochondria functionally utilize these various TRAP1 tetrameric states. This will involve the identification of unique subsets of client proteins for each state and mechanistic investigation of how each TRAP1 tetrameric state mediates client protein maturation. Finally, the observation of TRAP1 tetramers raises the possibility that oligomeric states could exist for other Hsp90 homologs under special cellular or organellar conditions.

## Acknowledgements

We thank members of the Agard Lab for helpful discussions. We gratefully thank David Bulkley, Eric Tse, Michael Braunfeld, and Glenn Gilbert from the W.M. Keck Foundation Advanced Microscopy Laboratory at the University of California, San Francisco (UCSF) for maintaining the EM facility and helping with data collection. Special thanks to Matt Harrington and Joshua Baker-LePain for computational support on the USCF Wynton cluster. This work was supported by funding from National Institutes of Health grants U54CA209891, S10OD020054, and S10OD021741. Y.L. was supported by a Howard Hughes Medical Institute-Helen Hay Whitney Foundation Postdoctoral Fellowship, an American Heart Association Postdoctoral Fellowship grant #18POST33990362, and the Program for Breakthrough Biomedical Research which is partially funded by the Sandler Foundation. M.S. was supported by 2018 AACR-Takeda Oncology Lymphoma Research Fellowship grant #18-40-38-SUN. D.A.A was supported by the Howard Hughes Medical Institute.

## Authors contributions

Y.L. and D.A.A. designed all the experiments. Y.L. and D.E. designed the construct, purified the proteins, and performed biochemical experiments. A.G.L. carried out the AUC experiment and the data analysis. Y.L. and M.S. collected and processed the cryo-EM data. Y.L. built the atomic models. D.A.A. supervised the project. Y.L. and D.A.A wrote the manuscript and all authors contributed to editing.

## Data availability

The electron density maps and atomic models have been deposited into the Electron Microscopy Data Bank (EMDB) and the Protein Data Bank (PDB). The accession codes are EMD-22918 and PDB-7KLU for TRAP1 tetramer, 22919 and PDB-7KLV for TRAP1 dimer.

