## Supporting Information for "Cryo-EM analysis of human mitochondrial Hsp90 in multiple tetrameric states"

This PDF file includes:

Methods
Figures S1 to S11
Tables S1
References

### METHODS

#### Protein cloning, expression, and purification

The expression constructs and purification procedures for human TRAP1 and $TRAP1_{SpyTag-SpyCatcher}$ are as same as previously described [1,2]. The $TRAP1_{SdhB}$ construct is derived $TRAP1_{SpyTag-SpyCatcher}$ by fusing a fragment of human SdhB (residue 29-160) to the C-terminus of $TRAP1_{SpyTag}$. A 24-residue linker including $(GGGGS)_3$ and HRV-3C cleavage sequence is inserted between SpyTag and SdhB. Expression was carried out in *E. coli* BL21(DE3)-RIL. Cells were grown in TB media (supplemented with 0.2 mM cysteine and 0.2 mM ferrous sulfate for SdhB fused $TRAP1_{SpyTag}$) at 37˚C to $OD_{600}$ of ~0.6 and then induced with 0.5 mM IPTG at 16 ˚C for 18 hr. SdhB fused $TRAP1_{SpyTag}$ and $TRAP1_{SpyCatcher}$ were expressed separately and then combined in roughly stoichiometric amounts after imidazole elution in Nickel-affinity chromatography and incubated overnight at 4˚C to form the heterodimer of $TRAP1_{SdhB}$ while dialyzing into a low-salt buffer (150 mM KCl and 20 mM Tris pH 8.0). The $TRAP1_{SdhB}$ heterodimer were further purified using Mono Q anion exchange and size exclusion chromatography before they were aliquoted and snap frozen with liquid $N_2$.

#### Blue native PAGE

Protein samples (TRAP1, yeast Hsc82, HtpG) with or without 100 mM DTT were separated on NativePAGE Novex 3-12% bis-Tris gels in NativePAGE running buffer along with protein size standards. The closed states of TRAP1 and Hsc82 were prepared by incubating the proteins ample with 1 mM AMP·PNP and 1 mM $MgCl_2$ at 30 °C for 1 hour. The cathode buffer contained 0.02% Coomassie brilliant blue G-250. Electrophoresis was performed at 4°C at 150 V for 2.5 h. Gels were then fixed by microwaving in 40% methanol and 10% acetic acid and incubating for 15 min and then destained by microwaving in 8% acetic acid and incubating until bands appeared on a clear background. Gels were then washed in water for 3 days.

#### Analytical ultracentrifugation

TRAP1 were dialyzed into 20 mM HEPES pH 8.0, 150 mM KCl, and 0.5 mM TCEP overnight at 4 °C. The samples were incubated at the appropriate temperature for 50 min in a pre-equilibrated rotor under vacuum. Runs were performed at 50,000 r.p.m. for 8–10 h in a Beckman XL/A analytical ultracentrifuge. Scans were collected at 260 or 280 nm with a radial step size of 0.003 cm at approximately 60-s intervals. SV analysis was done with SEDFIT/SEDPHAT(NIH) software.

#### Steady-state ATPase assay

The ATPase assay monitors phosphate release via a chromogenic substrate 7-methyl-6-thioguanosine (7-MESG) in the presence of an *E. coli* PNPase (purine nucleoside phosphorylase)[3]. In these assays, 0.14 mM 7-MESG and 2 µM PNPase was used per reaction. The change in absorbance of 7-MESG was measured at 355 nm. Initial rates from phosphate release assays were obtained from fitted slopes of a linear function, $y = mx + b$, to the most linear region of each kinetic trace.

#### Cryo-EM sample preparation

The closed state of all TRAP1 constructs was obtained by incubating the sample with 1 mM AMP·PNP and 1 mM $MgCl_2$ at 30 °C for 1 h. Cryo-EM grids were prepared with Vitrobot Mark IV (FEI, Thermo Fisher Scientific, Hillsboro), using 16°C and 100% humidity. 4 µL aliquots of samples were applied to glow discharged Quantifoil R1.2/1.3, 300-mesh copper holey carbon grids

(Quantifoil Micro Tools, GmbH, Großlöbichau, Germany), single blotted for 6-10 seconds with blot force 3, and plunge frozen in liquid ethane cooled by liquid nitrogen.

**Cryo-EM data collection**

Data were collected on a Titan Krios microscope (Thermo Fisher Scientific) operated at 300 kV with a K3 Summit direct electron detector (Gatan, Inc.). A GIF-BioQuantum energy filter with a slit width of 20 eV was used. Images were recorded using SerialEM[4] with defocus varied from -0.8 to -2.2 μm for a total dose of 66 $e^-/Å^2$. A super-resolution pixel size of 0.425 Å was used and each image was dose-fractionated to 100 frames (0.06 s each, total exposure of 6 s) with a dose rate of 6.6 $e^-/Å^2/s$. A summary of the data collection parameter was provided in Supplementary Table 1.

**Image processing**

Image stacks were motion-corrected and summed using MotionCor2[5], resulting in Fourier-cropped summed images which are binned by 2. CTFFIND4 was used to estimate defocus parameters for all the images[6]. Initial particle picking was carried out using Gautomatch without a template to generate the 2D class averages, which were then used as templates for a second-round particle picking on micrographs with 25 Å low-pass filtering. Relion 3.1[7,8] was used for all the following steps. One round of reference-free 2D classification were performed for 25 iterations each with images binned by 4. Good particles were picked from 2D averages, extracted as images binned by 2, and subjected to two rounds 3D classification within Relion. An initial model was generated from cryo-EM map of human TRAP1 (EMD-22174 and PDB 6XG6)[9] and low-pass filtered to 30 Å. Classes corresponding to TRAP1 dimer and tetramer were re-extracted without binning, and subjected to 3D auto-refinement. All refined maps were post-processed and sharpened by an automatically estimated B-factor. All resolutions were estimated by applying a soft mask around the protein density and the gold-standard Fourier shell correlation (FSC) = 0.143 criterion.

**Model building, refinement and validation**

The initial model of TRAP1 dimer and tetramer was derived from the cryo-EM structure of human TRAP1 bound with free SdhB (EMD-22812 and PDB 7KCL, to be released)[10]. The docked models were refined in real space against the cryo-EM maps using real space refinement in PHENIX[11] with secondary structure restraints, followed by iterative rounds of manual and automated refinement in Coot[12] and PHENIX[11], respectively. The final models and their fitness into the maps were visually inspected and geometry was further evaluated using MolProbity[13]. Cryo-EM data collection, refinement and validation statistics are summarized in Supplementary Table 1. Figures depicting the structures were prepared in ChimeraX[14] or VMD[15].

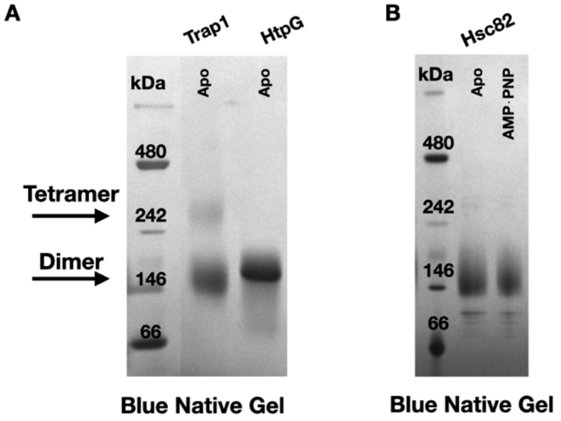

**Figure S1.** Blue native PAGE analysis of Hsp90s. (A) TRAP1 forms both tetramer and dimer, whereas HtpG only forms dimer. (2) Yeast cytosolic Hsc92 only forms dimer with or without AMP·PNP.

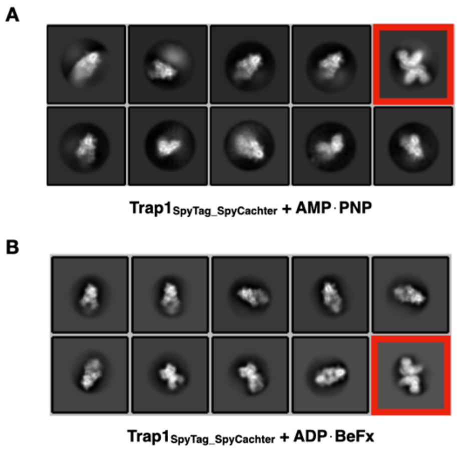

**Figure S2.** Representative 2D-averaged classes of TRAP1 heterodimer covalently linked with SpyTag and SpyCatcher (TRAP1$_{SpyTag\text{-}SpyCatcher}$) in the presence of (A) AMP·PNP or (B) ADP·BeF$_x$. The "butterfly" tetramer conformation are boxed in red squares.

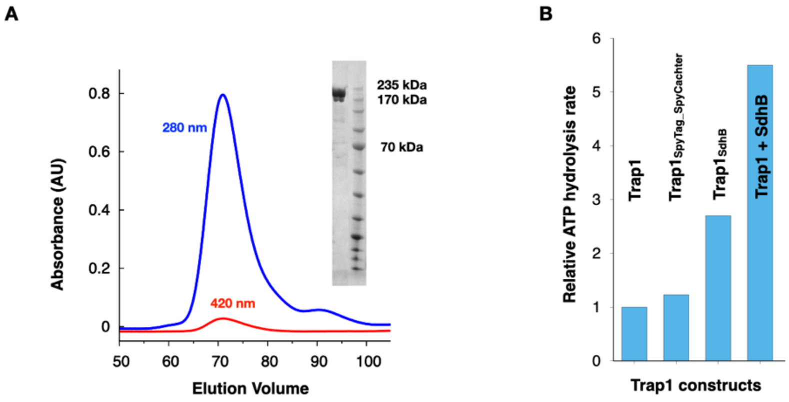

**Figure S3.** Characterization of TRAP1 heterodimer fused with SdhB (TRAP1$_{SdhB}$). (A) Gel filtration profile and SDS-PAGE for TRAP1$_{SdhB}$. (B) Relative ATPase activity of various TRAP1 constructs.

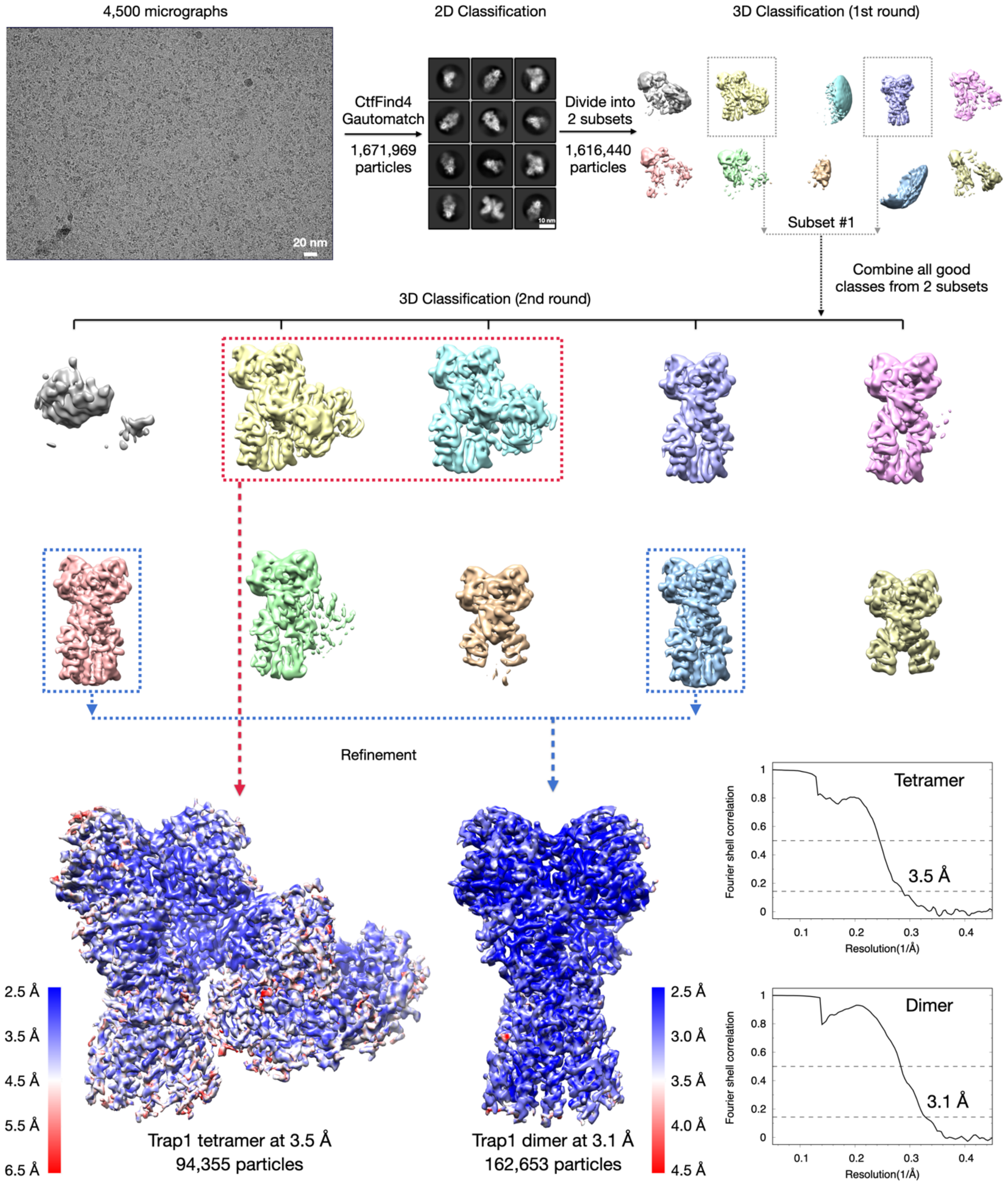

**Figure S4.** Cryo-EM workflow for image analysis. Local resolution maps and gold-standard Fourier shell correlation (FSC) curves for cryo-EM structures of TRAP1 tetramer and dimer wer shown at the bottom.

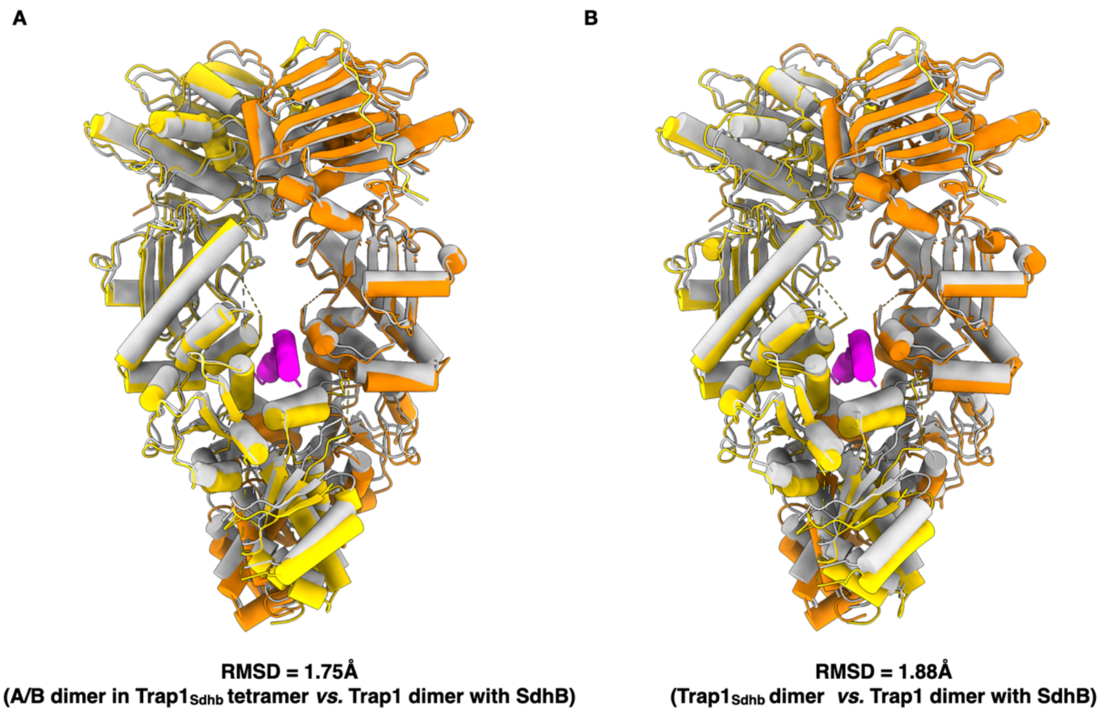

**Figure S5.** Structural comparison between TRAP1 bound with free SdhB (gray for TRAP1 and magenta for SdhB) and with fused SdhB (orange and yellow for TRAP1 protomer A and B, respectively). (A) The structural alignment between A/B dimer in the TRAP1 tetramer and TRAP1 bound with free SdhB. (B) The structural alignment between in the TRAP1 dimer with fused Sdhb and TRAP1 bound with free SdhB.

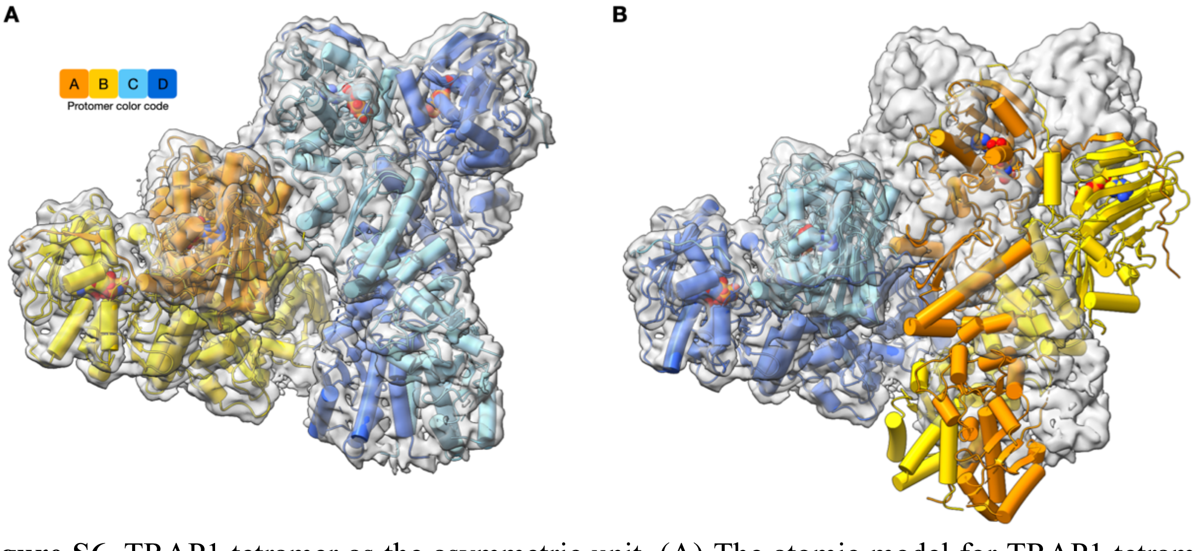

**Figure S6.** TRAP1 tetramer as the asymmetric unit. (A) The atomic model for TRAP1 tetramer built into the cryo-EM density. (B) The mismatch between the atomic model of A/B dimer and the cryo-EM density of C/D dimer in the TRAP1 tetramer after aligning the atomic model of C/D dimer into the cryo-EM density of A/B dimer.

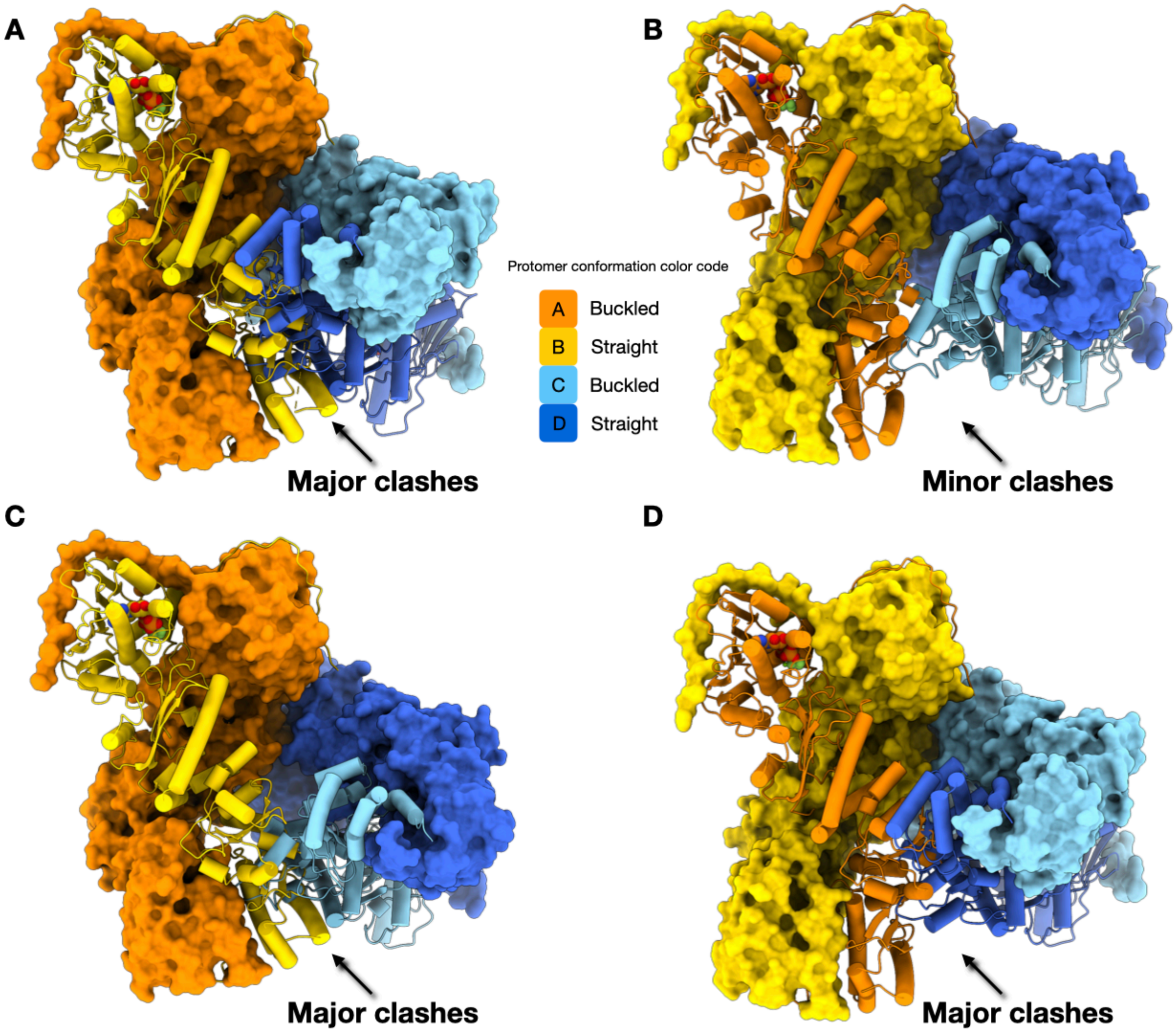

**Figure S7.** Docking of the asymmetric closed state of TRAP1 dimers into the tetrameric state by preserving the MD dimer-dimer interface I. (A) MD dimer-dimer interface is formed between the buckled protomers A and C. A major clash is observed between straight protomers B and D. (B) MD dimer-dimer interface is formed between straight protomers B and D. A minor clash is observed between the buckled protomers A and C. (C) MD dimer-dimer interface is formed between the buckled protomer A and straight protomer D. A major clash is observed between straight protomer B and buckled protomer C. (B) MD dimer-dimer interface is formed between straight protomer B and buckled protomer C. A major clash is observed between the buckled protomers A and straight D.

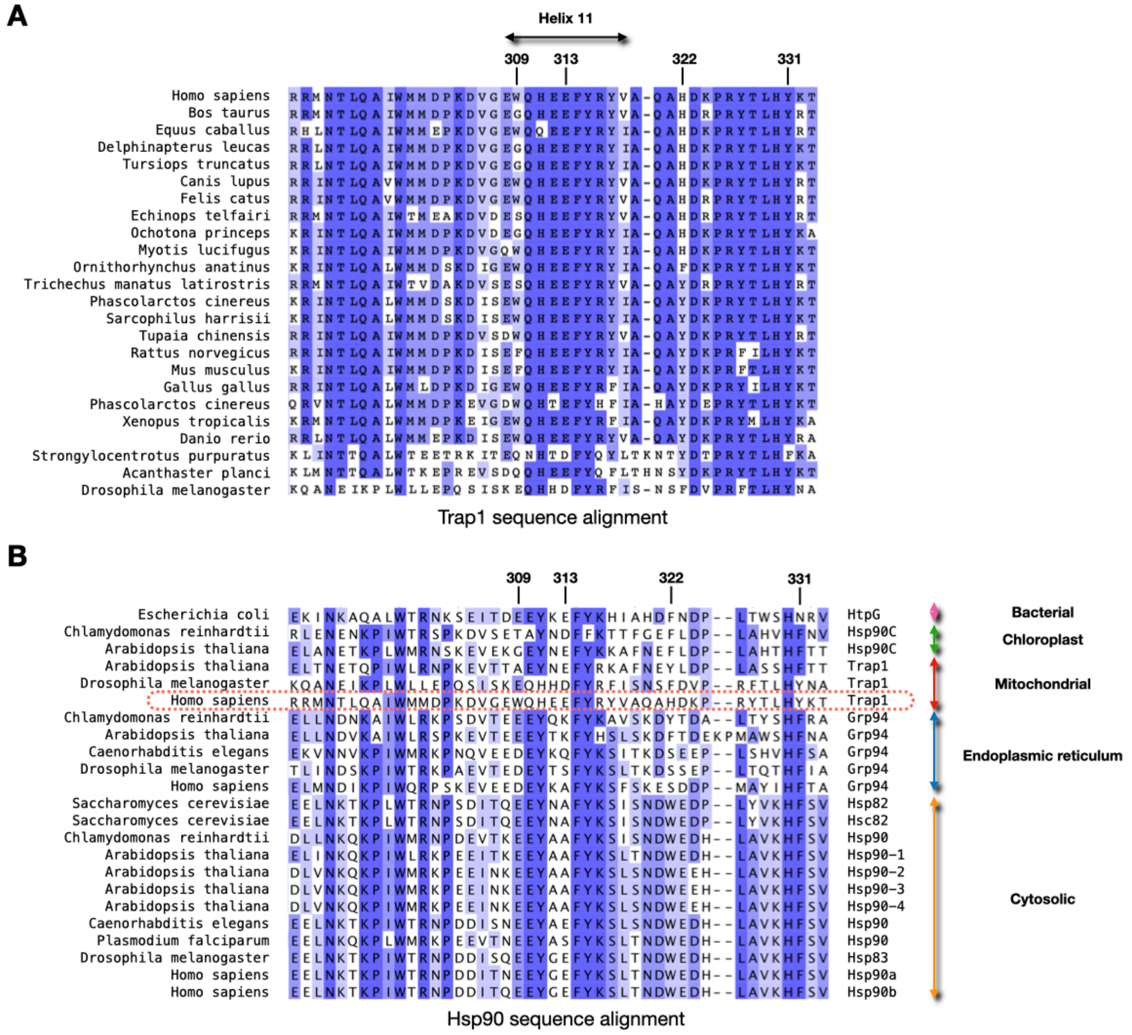

**Figure S8.** Sequence alignment of (A) TRAP1 and (B) all Hsp90 homologs around helix 11 region.

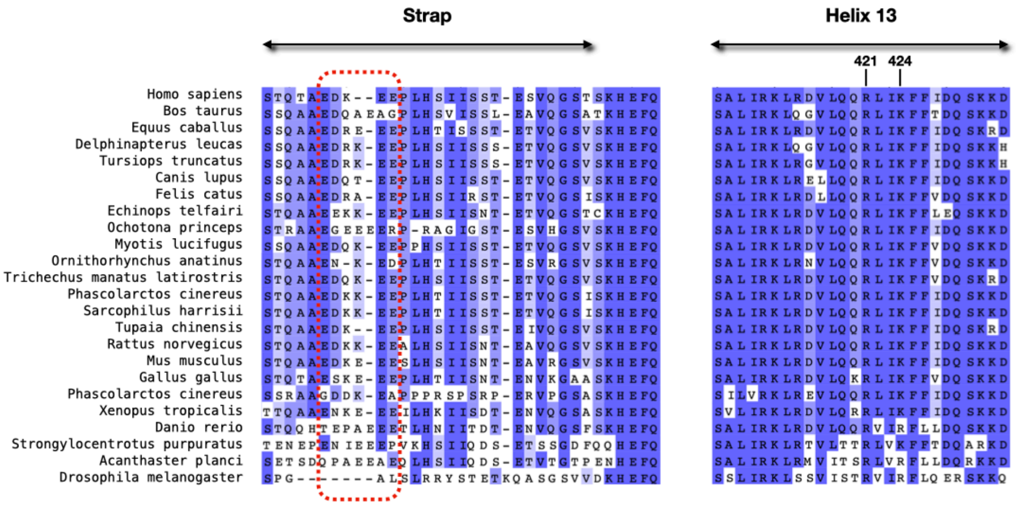

**Figure S9.** Sequence alignment of TRAP1 around strap and helix 13 region.

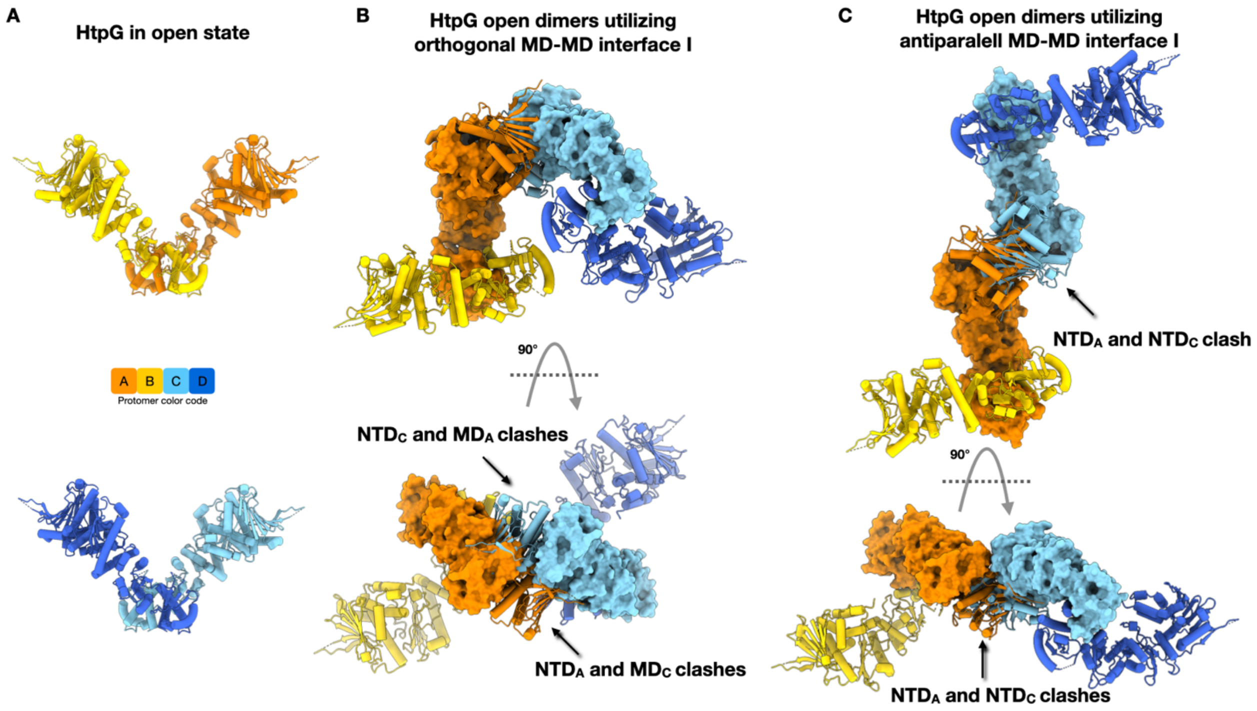

**Figure S10.** Docking of the open state of HtpG dimers into the tetrameric state based only the MD dimer-dimer interface I. (A) The open state of HtpG dimers. (B) The orthogonal MD dimer-dimer interface is formed between the protomers A and C. Major clashes are observed between NTD from protomer A ($NTD_A$) and MD from protomer C ($MD_C$), as well as between NTD from protomer C ($NTD_C$) and MD from protomer A ($MD_A$). (C) The antiparallel MD dimer-dimer interface is formed between the protomers A and C. Major clashes are observed between NTDs from protomer A and C.

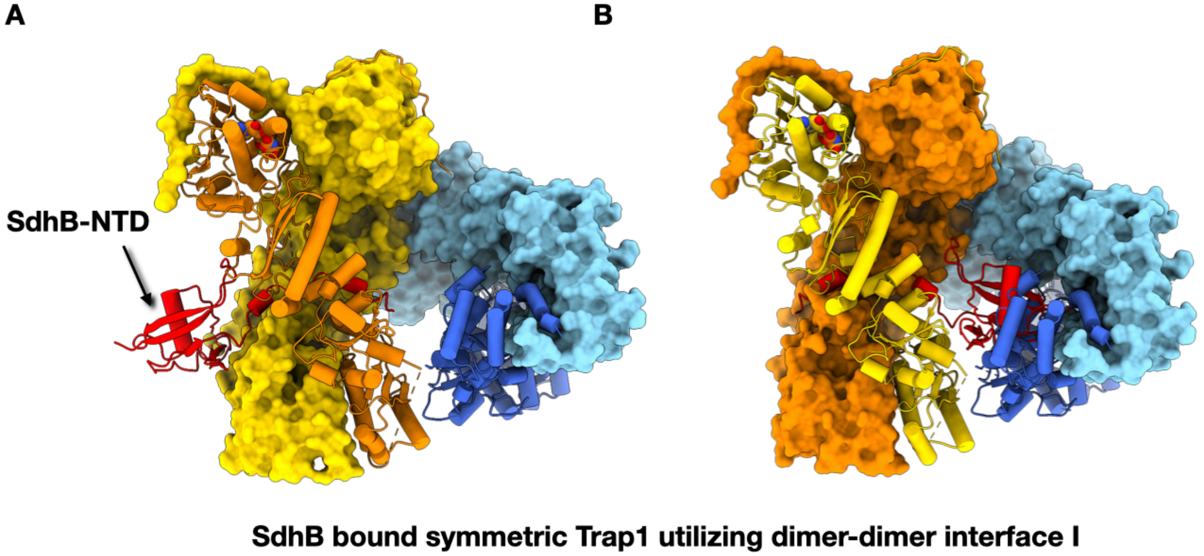

**Figure S11.** Docking of TRAP1 dimer bound with free SdhB into the TRAP1 tetrameric state. (A) A folded SdhB-NTD is compatible with TRAP1 tetramer if bound to the outer face. (A) The SdhB-NTD is incompatible with TRAP1 tetramer if bound between the dimers.

| | #1 TRAP1_tetramer (EMD-22918) (PDB 7KLU) | #2 TRAP1_dimer (EMD-22919) (PDB 7KLV) |
|---|---|---|
| **Data collection and processing** | | |
| Microscope | FEI Titan Krios | FEI Titan Krios |
| Camera | Gatan K3 Summit | Gatan K3 Summit |
| Energy Filter | 20 eV slit | 20 eV slit |
| Magnification | 105,000x | 105,000x |
| Voltage (kV) | 300 | 300 |
| Electron exposure (e–/Å$^2$) | 66 | 66 |
| Defocus range (μm) | -0.8 to -2.3 | -0.8 to -2.3 |
| Pixel size (Å) | 0.849 | 0.849 |
| Symmetry imposed | C1 | C1 |
| Micrographs (no.) | 4,500 | 4,500 |
| Initial particle images (no.) | 1,671,969 | 1,671,969 |
| Final particle images (no.) | 94,355 | 162,653 |
| Map resolution (Å) | 3.5 | 3.1 |
| FSC threshold 0.143 | | |
| Map resolution range (Å) | 3.0-6.0 | 2.5-4.0 |
| **Refinement** | | |
| Initial model used (PDB code) | 6XG6 | 6XG6 |
| Model resolution (Å) | 3.7 | 3.4 |
| FSC threshold 0.5 | | |
| Map sharpening *B* factor (Å$^2$) | -60 | -80 |
| Model composition | | |
| Non-hydrogen atoms | 19742 | 9871 |
| Protein residues | 2452 | 1226 |
| Ligands | 2 | 2 |
| Ions | 4 | 4 |
| *B* factors (Å$^2$) | | |
| Protein | 77.43 | 84.68 |
| Ligand | 50.08 | 54.54 |
| R.m.s. deviations | | |
| Bond lengths (Å) | 0.002 | 0.005 |
| Bond angles (°) | 0.522 | 0.639 |
| Validation | | |
| MolProbity score | 1.63 | 1.77 |
| Clashscore | 5.94 | 7.53 |
| Poor rotamers (%) | 0.00 | 0.00 |
| Ramachandran plot | | |
| Favored (%) | 95.66 | 94.79 |
| Allowed (%) | 4.34 | 5.21 |
| Disallowed (%) | 0.00 | 0.00 |

**Table S1.** Cryo-EM data collection, refinement and validation statistics
